# CD32 allows capturing blood cells emergence in slow motion during human embryonic development

**DOI:** 10.1101/2023.03.23.530597

**Authors:** Rebecca Scarfò, Monah Abou Alezz, Mahassen El Khoury, Amélie Gersch, Zhong-Yin Li, Stephanie A. Luff, Sara Valsoni, Sara Cascione, Emma Didelon, Ivan Merelli, Jean-Noël Freund, Christopher M. Sturgeon, Manuela Tavian, Andrea Ditadi

## Abstract

During development, in the embryo proper blood cells emerge from a subset of specialized endothelial cells, named hemogenic endothelial cells (HECs), via a process known as endothelial-to-hematopoietic transition (EHT) driven by time-specific Notch signaling activation^1^. HECs represent an elusive cell population as they are rare and transient, rapidly generating blood cells, and specific markers are lacking. Therefore, it remains unclear how and when the hematopoietic fate is specified and how blood cell emergence is molecularly regulated. Notably, thorough characterization of this process is essential to guide the generation of therapeutic blood products *in vitro* from human pluripotent stem cells (hPSCs). To identify specific human HEC markers, we performed transcriptomic analysis of 28-32-day human embryos, a developmental stage characterized by active hematopoiesis. We observed that the expression of *FCGR2B*, encoding for the Fc receptor CD32, is highly enriched in the ACE^+^CD34^+^ endothelial cell population that contains HECs. Functional *ex vivo* analyses confirmed that multilineage hematopoietic potential is highly enriched in CD32^+^ endothelial cells isolated from human embryos. In addition, clonal analysis revealed that 90% of CD32^+^ hPSC-derived endothelial cells are *bona fide* HECs. We leveraged this specificity to study how HECs commit to the blood fate. Remarkably, our analyses indicated that HECs progress through different states culminating with the one identified by CD32 expression. Indeed CD32^+^ HECs no longer require Notch to generate hematopoietic progeny and display full commitment to hematopoiesis even before the expression of hematopoietic markers. These findings provide a precise method for isolating HECs primed to the blood fate from human embryos and hPSC cultures, thus allowing the efficient generation of hematopoietic cells *in vitro*.

## Main

During embryonic development, cells continuously alter their characteristics to specify new fates and generate the entire spectrum of cell types and tissues. Sometimes cell differentiation involves abrupt lineage conversions as in the case of embryonic hematopoiesis. In fact, various experimental models have informed us that, during development, hematopoietic cells are produced from hemogenic endothelial cells (HECs), a specialized subpopulation of embryonic endothelium^2–11^. HECs are thought to generate blood cells via an endothelial-to-hematopoietic transition (EHT), which involves considerable transcriptional and morphological changes leading to the identity switch from endothelial cell to blood^2, 3, 5–8, 12^. HECs represent a central element of the distinct hematopoietic developmental programs. In fact, HECs can be found in the different anatomical locations where hematopoiesis is observed, including the yolk sac (YS), where lineage-restricted hematopoietic progenitors are generated first, as well as the aorta-gonad-mesonephros (AGM), the site of hematopoietic stem cells (HSC) emergence, where hematopoiesis occurs in a Notch-dependent manner^13–15^. As HECs represent a rare and transient population rapidly generating hematopoietic output, they have been difficult to characterize and therefore little is known about how the emergence of the hematopoietic lineage is orchestrated.

Traditionally, HECs have been isolated and characterized in animal models, using reporters under the control of the regulatory elements of the transcription factors that drive blood cell emergence, such as *Runx1* and *Gfi1*^16–18^, a strategy that cannot be used to study HECs in human embryos. Our previous data suggested angiotensin-converting enzyme (ACE, also known as CD143) as a potential marker of HSCs and their endothelial precursors in human embryo^19, 20^. Recently, transcriptomic analyses have allowed the identification of putative HEC markers in both murine and human embryos, including ACE as well as CXCR4 and CD44^21–24^, but these also enrich for arterial endothelial cells, anatomically associated with HECs^25–27^. This hinders the specific characterization of the unique endothelial population that generate blood cells and, consequently, the design of accurate protocols for the derivation of therapeutically relevant hematopoietic cells from hPSCs.

To overcome these limitations and identify broadly applicable cell-surface markers specific for human HECs both *in vivo* and *in vitro*, we performed transcriptomic analysis of endothelial populations displaying hematopoietic potential in the human embryo based on ACE expression. Here we report that the Fc receptor CD32 is expressed on human embryonic endothelial cells with robust hemogenic potential. Likewise, CD32 allows for the identification of hPSC-derived HECs with higher specificity than other reported HEC markers. This provided the unprecedented opportunity to study how and when HECs initiate the hematopoietic program. We show that HECs transit through different states and their hematopoietic commitment occurs before the expression of hematopoietic markers and cell cycle genes. In particular, CD32 marks a specific subpopulation of HECs which no longer require Notch activation and that are fully committed to hematopoiesis. The ability to capture this rare and rapid embryonic process allows redefining how blood cells emerge in the embryo and will enable the efficient generation of hematopoietic cells *in vitro* for therapies.

### ACE expression distinguishes endothelial cells with distinct gene expression programs in human embryos

Using spatio-temporal analyses, we have previously shown that within the AGM region of the human embryo, hematopoietic clusters firstly emerge in the dorsal aorta (DA) at 27 days post-fertilization (dpf; Carnegie Stage (CS) 12)^28^. Human hematopoietic clusters are located on the ventral side of the aortic endothelium and express CD34, a surface marker that they share with the surrounding endothelial cells^28, 29^. Intra-aortic CD34^+^ hematopoietic clusters also express ACE, another marker that identifies cells with hematopoietic potential in the developing human embryo^19, 20^. As ACE expression is also observed in endothelial cells within different hematopoietic sites during human development, including the DA^19, 20^, we further analyzed its expression pattern in the AGM region in parallel with the expression of the transcription factor RUNX1, which regulates the emergence of blood cells in the embryo^16, 17^. Immunofluorescence analysis on human embryonic sections revealed that at 23 dpf (CS10), ACE^+^ cells can be found in the mesenchyme surrounding the DA, but not in the aortic wall (Fig. 1a). At this stage RUNX1 expression cannot be observed in the human AGM region. However, at 27 dpf (CS12), when the first hematopoietic clusters emerge inside the DA, ACE and RUNX1 expression colocalized in the endothelial cells lining the ventral site of the DA (Fig. 1a). As such, ACE expression emerges as a potential human HEC marker *in vivo*, similarly to what is described for murine HECs^23^.

**Figure 1.**
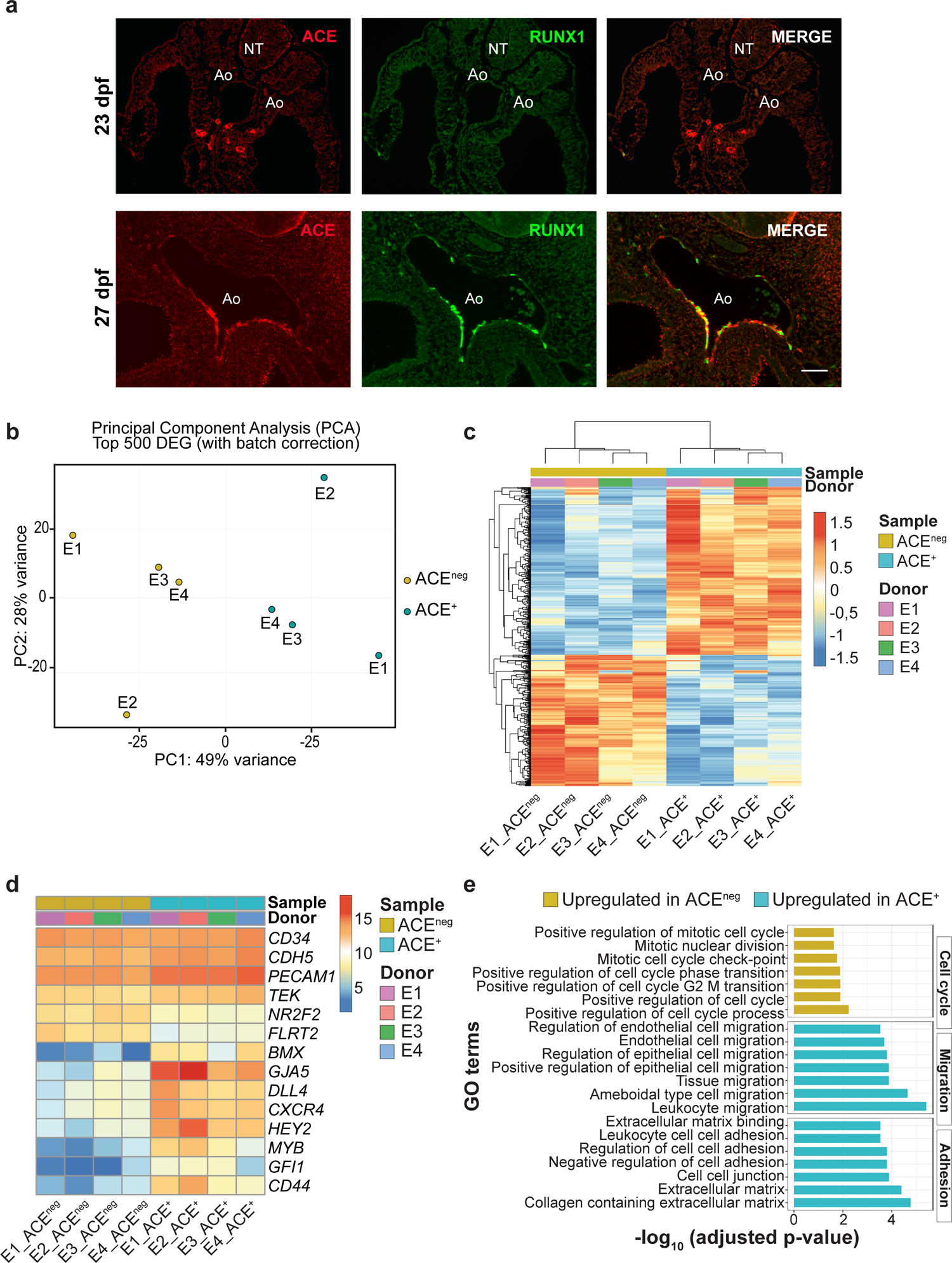
Human embryonic ACE+ endothelial cells express arterial and hemogenic markers. **a**) Transverse section of the AGM region of a 23 dpf (CS10, upper panels, *n*=2 independent) and a 27 dpf (CS12, lower panels, *n*=3 independent) human embryo, immunostained with ACE (left panels, in red), RUNX1 (middle panels, in green) and merge (right panels). Ao: aorta; NT: neural tube. Scale bar: 50 μm; **b**) Principal component analysis (PCA) of the top 500 DEGs within indicated human embryonic populations (ACE^neg^ in beige and ACE^+^ in light blue) isolated from four CS12-13 human embryos; **c**) Heatmap visualizing the DEGs within indicated human embryonic populations (ACE^neg^ in beige and ACE^+^ in light blue) isolated from four CS12-13 human embryos (referred to “donor”: E1, E2, E3, E4). Gene counts were corrected for donor. The rlog gene expression values are shown in rows and tiles referring to DEGs are coloured according to up-(red) or down-(blue) regulation; **d**) Heatmap visualizing the expression of selected pan-endothelial (*CD34, CDH5, PECAM, TEK*), vein-specific (*NR2F2, FLRT2*), arterial-specific (*GJA5, DLL4, CXCR4, HEY2*) and hemogenic (MYB, GFI1, CD44) genes within ACE^neg^ (beige) or ACE^+^ (light blue) populations isolated from the four CS12-13 human embryos E1, E2, E3 and E4 (donor). The rlog gene expression values are shown in rows. Tiles referring to the DEGs are colored according to up- (red) or down- (blue) regulation; **e**) Barplot showing significantly enriched Gene Ontology (GO) terms (adjusted p-value<0.05) using ORA on DEGs from the comparison between ACE^+^ (light blue) versus ACE^neg^ (beige) samples. The barplot shows enriched terms grouped by custom categories: cell cycle (upregulated in ACE^neg^), migration and adhesion (upregulated in ACE^+^).

We then tested whether ACE can also distinguish subsets of endothelial cells *in vitro* and track hematopoietic potential in hPSC hematopoietic cultures. For this, we differentiated the human embryonic stem cell (hESC) line H1 using our protocol that specifies hPSCs into WNT-dependent (WNTd) NOTCH-dependent multipotent *HOXA*^+^ HECs, indicative of intra-embryonic AGM-like hematopoiesis^27, 30–32^. In this setting, we observed that ACE is expressed by virtually all day 8 CD34^+^ cells, including 90±1% of CD34^+^CD43^neg^CD73^neg^CD184^neg^ cells that comprise HECs in day 8 WNTd hPSC hematopoietic cultures (Extended data Fig. 1a, b)^27^. Since HECs represent only:::2% of WNTd CD34^+^CD43^neg^CD73^neg^CD184^neg^ cells^27^, we concluded that ACE expression is unable to distinguish HECs from other vascular endothelial cells in these hPSC differentiating cultures.

Having established that ACE expression segregates with the hemogenic transcription factor RUNX1 in DA endothelial cells of the human embryo, we used this surface marker to better characterize embryonic endothelial cells with hematopoietic potential and identify markers that track with HECs both *in vivo* and *in vitro*. For this, we isolated CD34^+^CD45^neg^ cells based on their ACE expression from the AGM region of 4 human embryos (E1, E2, E3, E4) staged between 28 and 32 dpf (CS12-13; Extended Data Fig. 1c) and performed whole-transcriptomic analysis using bulk RNA sequencing (RNAseq). Principal component analysis (PCA) and unsupervised hierarchical clustering by K-Means highlighted a clear segregation of CD34^+^CD45^neg^ACE^+^ and CD34^+^CD45^neg^ACE^neg^ (herein ACE^+^ and ACE^neg^, respectively) transcriptomes (Fig. 1b, c, Extended Data Fig. 1d, e), with 785 differentially expressed genes (DEGs), of which 440 genes upregulated and 345 downregulated in ACE^+^ (Fig. 1c and Supplementary Table 1a). At the population level, both cell fractions expressed high levels of endothelial genes (*CD34*, *CDH5*, *PECAM1*, *TEK*) while genes associated with venous fate (*NR2F2*, *FLRT2*) of endothelial cells had a lower expression in ACE^+^ cells in comparison to ACE^neg^ cells. In contrast, ACE^+^ cells showed a higher expression of genes classically associated with arterial cells (*BMX, GJA5*, *DLL4*, *CXCR4*, *HEY2*) and HECs (*MYB*, *GFI1*, *CD44*; Fig. 1d). Over representation analysis (ORA) and gene-set enrichment analysis (GSEA) revealed that ACE^+^ cells are positively associated with cell migration- and cell adhesion-related GO terms, in line with the remodeling occurring during the emergence of blood cells from HECs^33^ (Fig. 1e and Supplementary Table 1b-e). Conversely, ACE^+^ cells are negatively associated with several cell cycle-related GO-terms, suggesting that at a population level they are undergoing an active arterialization process which requires cell growth suppression^34^. These results indicated that in the aortic endothelium of 5-week human embryos, ACE expression identifies cells showing an arterial gene signature as well as cells expressing genes pivotal for hematopoietic development.

### CD32 is expressed in HECs in human embryonic hematopoietic regions

To identify putative markers for the isolation of HECs, we focused our attention on DEGs coding for cell surface proteins showing higher expression in ACE^+^ cells sorted from human embryos (Fig. 2a, Extended Data Fig. 2a and Supplementary Table 1f). We found that *FCGR2B*, which encodes for an isoform of the Fc receptor CD32, ranks among the top 10 cell-surface genes whose expression is enriched in ACE^+^ cells. Given that CD32 is a marker of other specialized endothelial cells (such as liver sinusoidal endothelial cells^35^, placenta villi^36^ and dermal microvasculature^37^) and that Fc receptors are expressed together with endothelial markers in a subset of YS-derived hematopoietic progenitors^38^, we focused our attention on this gene. We performed immunohistochemical analysis on human embryo sections between 26-30 dpf (CS12-13), which showed that CD32 is expressed together with CD34 in the aortic endothelial cells. CD32^+^ endothelial cells localized close to the bifurcation with the vitelline artery (VA) in the AGM region, a site known to contain a high frequency of hematopoietic clusters, and therefore potentially of HECs, at this stage (Fig. 2b)^28^. Remarkably, in the DA CD32 marks both the intra-aortic hematopoietic clusters bordering on the ventral site and the underlying endothelial cells. Immunofluorescence analysis on consecutive sections showed that CD32^+^ endothelial cells in the DA co-express CD34, ACE and RUNX1 (Fig. 2c). In the CS12 human embryo, CD32 is also expressed together with CD34 in endothelial cells surrounding the hemogenic regions of the YS (Fig. 2d). This specific expression pattern at active hemogenic embryonic sites led us to functionally test whether CD32 identifies HECs. Thus, we isolated by FACS the CD32^+^ and CD32^neg^ fractions of the CD34^+^CD43^neg^CD45^neg^ endothelial cell population containing HECs in the AGM and YS regions dissected from two CS13 embryos (E5, E6) and assessed their potential to generate hematopoietic progenitors *ex vivo* (Fig. 2e, and Extended Data Fig. 2b, c, d). After 7 days of culture on OP9DLL1 stroma, the CD32^+^ endothelial fraction from both the AGM and YS regions of both embryos consistently generated more clonogenic progenitors with erythro-myeloid potential than the CD32^neg^ cells (Fig. 2f). Altogether, these results demonstrate that CD32 can identify a subset of endothelial cells with robust hematopoietic potential in the human embryo.

**Figure 2.**
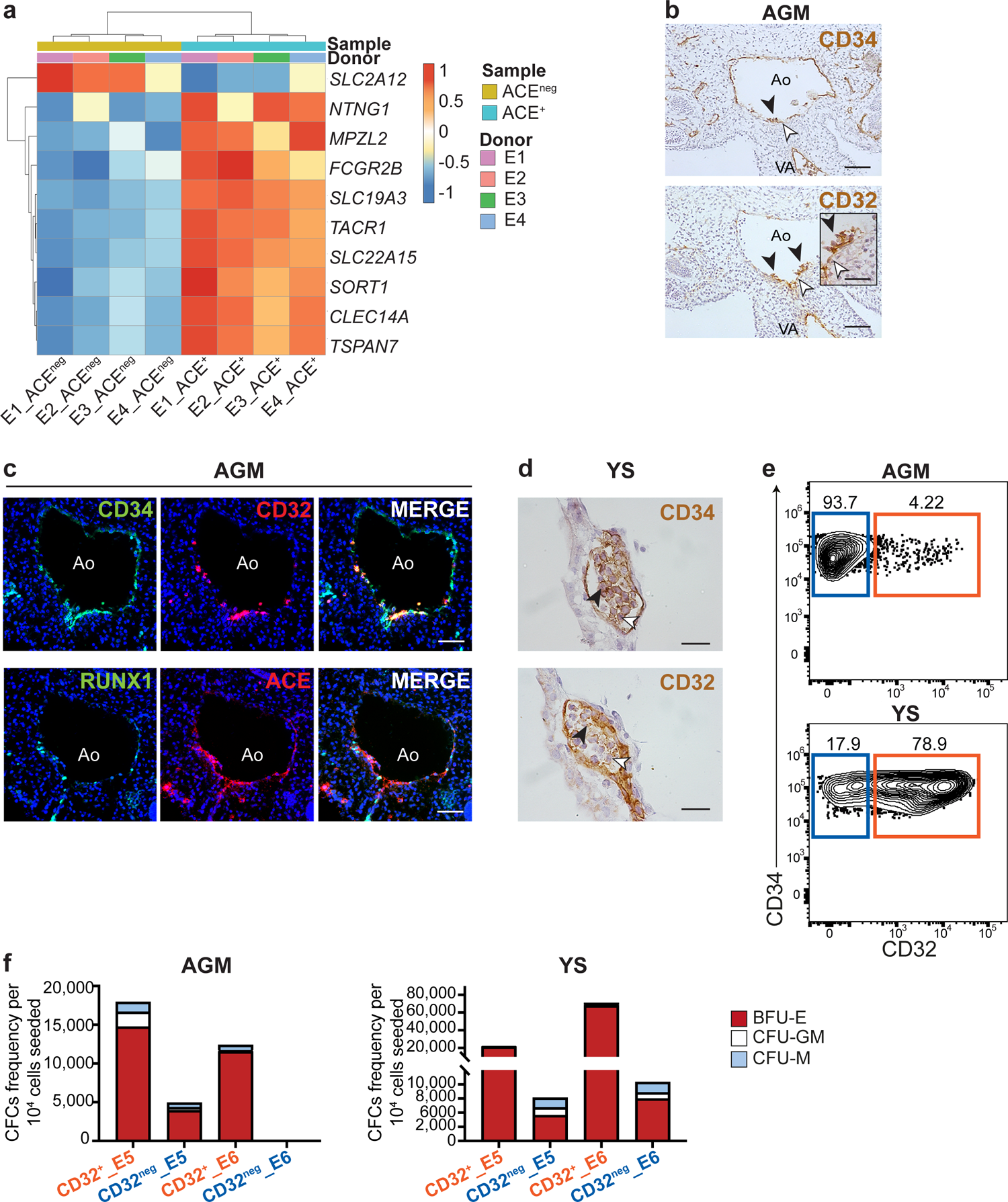
CD32 is expressed in AGM and YS HECs during human embryonic development. **a**) Heatmap visualizing the top 10 differentially expressed cell surface genes within indicated human embryonic populations (ACE^+^ in light blue and ACE^neg^ in beige). Cells are derived from the four CS12-13 human embryos E1, E2, E3 and E4 (donor). The rlog gene expression values are shown in rows. Tiles referring to differentially expressed cell surface genes are colored according to up-(red) or down- (blue) regulation; **b**) CD34 (upper panel) and CD32 (lower panel) expression by immunohistochemistry of consecutive sections of the AGM of a 29 dpf (CS13) human embryo (*n*=5 independent). Inset shows a high magnification of the hematopoietic clusters (black arrowhead) and surrounding endothelial cells (white arrowhead) in the aorta immunostained by CD32. Ao: aorta; VA: vitelline artery. Scale bars: 50 μm and 100 μm in the inset; **c**) Transverse consecutive sections of the AGM region of a 29 dpf (CS13) human embryo, immunostained with: upper panel from left to right, CD34 (green), CD32 (red) and merge; lower panel from left to right, RUNX1 (green), ACE (red) and merge; Ao: aorta. Scale bar: 50 μm; **d**) CD34 (upper panel) and CD32 (lower panel) expression by immunohistochemistry of consecutive sections of the YS of a 26 dpf (CS12) human embryo (*n*=5 independent); **e**) Representative flow cytometric analysis of CD32^+^ (orange) and CD32^neg^ (blue) cell populations within AGM and YS of CS13 human embryo. *n*=2, independent. Gated on SSC/FSC/Live/CD34^+^CD43^neg^CD45^neg^ as shown in Extended Data Fig. 2c, d; **f**) Quantification of erythro-myeloid CFC potential from CD32^+^ and CD32^neg^ populations isolated from the AGM (left panel) and YS (right panel) of the two CS13 human embryos (E5, E6) and cultured on OP9DLL1; *n*=2, independent. BFU-E: burst forming unit erythroid; CFU-GM: colony forming unit granulocyte macrophage, CFU-M: colony forming unit macrophage.

### CD32 defines intra-embryonic HECs with multilineage potential in hPSC hematopoietic cultures

Based on our results obtained with the human embryos, we next investigated whether CD32 can be a reliable marker for isolating HECs from hPSC differentiating cultures. We monitored its expression in WNTd intra-embryonic-like *HOXA*^+^ HECs^27, 30, 32, 39^ (Extended Data Fig. 3a). Using these conditions, we have previously shown that HECs displaying multipotent hematopoietic potential are contained within the CD34^+^CD43^neg^CD73^neg^CD184^neg^DLL4^neg^ cell fraction of day 8 hPSC differentiating cultures^27^. Remarkably, CD32 expression identified only a small subset of the HEC-containing CD34^+^CD43^neg^CD73^neg^CD184^neg^DLL4^neg^ cell population, which was all ACE^+^ (Fig. 3a, b and Extended Data Fig. 3b, c). In parallel, using the H9 hESC reporter line to evaluate the expression of *RUNX1C*, a RUNX1 isoform regulating WNTd hematopoieis^27, 30^, we observed that CD32 expression in day 8 WNTd CD34^+^CD43^neg^CD73^neg^CD184^neg^DLL4^neg^ cells demarcates endothelial cells that do not express *RUNX1C*-EGFP and thus might identify a population of HECs that have not initiated the hematopoietic program yet (Fig. 3c, d). To monitor whether this population can generate blood cells, we isolated CD32^+^ from CD34^+^CD43^neg^CD73^neg^CD184^neg^DLL4^neg^ day 8 WNTd cells and cultured them on Matrigel in the presence of hematopoietic cytokines known to promote and sustain hematopoietic differentiation (HEC culture), as we previously described^27^. Under these conditions, the cells formed an adhesive monolayer that generated hematopoietic progeny as demonstrated by the presence of round cells and of a population of *RUNX1C*-EGFP^+^ and CD45^+^ cells five days later (Fig. 3e, f). As such, the CD32^+^ fraction displays the defining behavior of *bona fide* HECs.

**Figure 3.**
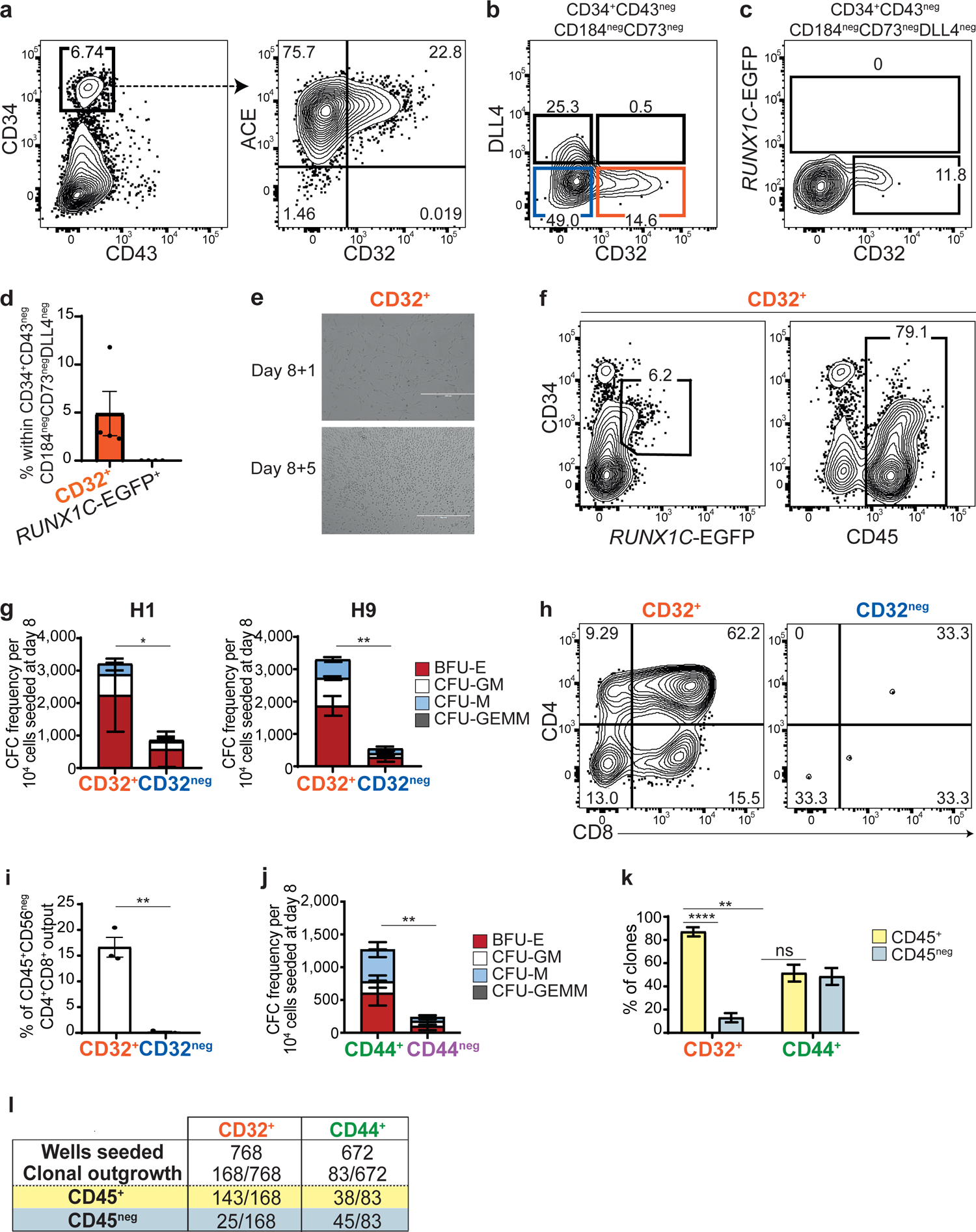
CD32 is expressed in WNT-dependent hPSC-derived HECs. **a**) Representative flow cytometric analysis of day 8 WNTd hPSC-derived hematopoietic cultures. Left panel: CD43 and CD34 expression gated on SSC/FSC/Live. Right panel: CD32 and ACE expression gated on SSC/FSC/Live/CD34^+^CD43^neg^ cells. H1 hESCs, *n*=3, independent; **b**) Representative flow cytometric analysis of CD32 and DLL4 expression within CD34^+^CD43^neg^CD184^neg^CD73^neg^ cells at day 8 of WNTd hPSC-derived hematopoietic cultures. CD32^+^ cells are shown in orange and CD32^neg^ cells in blue. Gated on SSC/FSC/Live/CD34^+^CD43^neg^CD184^neg^CD73^neg^. H1 hESCs, *n*=7, independent; **c**) Representative flow cytometric analysis showing CD32 and *RUNX1C*-EGFP expression in day 8 WNTd hPSC-derived hematopoietic cultures. Gated on SSC/FSC/Live/CD34^+^CD43^neg^CD184^neg^CD73^neg^DLL4^neg^. H9 hESCs, *n*=4, independent; **d**) Bar plot showing the frequency of CD32 and *RUNX1C*-EGFP expression within day 8 WNTd CD34^+^CD43^neg^CD184^neg^CD73^neg^DLL4^neg^ H9 hESCs in *n*=4 independent experiments, mean±SEM, as shown in Fig. 3c. Gated on SSC/FSC/Live/CD34^+^CD43^neg^CD184^neg^CD73^neg^DLL4^neg^; **e**) Photo-micrographs of CD32^+^ cells isolated at day 8 of WNTd hPSC-derived hematopoietic culture and cultured for 1 (day 8+1, upper panel) or 5 (day 8+5, bottom panel) days in the presence of pro-hematopoietic cytokines (HEC culture). At day 8+1, CD32^+^ cells appear adherent while round cells in suspension are present at day 8+5. H9 hESCs, *n*=3, independent; **f**) Representative flow cytometric analysis of endothelial (CD34^+^CD45^neg^) and hematopoietic cells (CD45^+^, *RUNX1C*-EGFP^+^) derived from CD32^+^ cells isolated at day 8 of WNTd hPSC-derived hematopoietic culture and analyzed after 5 days of culture in the presence of pro-hematopoietic cytokines (HEC culture). H9 hESCs, *n*=3, independent; **g**) Quantification of erythro-myeloid CFC potential of CD32^+^ and CD32^neg^ populations isolated from day 8 WNTd hPSC-derived hematopoietic cultures and cultured on OP9DLL1, H1 hESCs (*n*=4), on the left, and H9 hESCs (*n*=3), on the right. One-tail paired Student’s t-test, nonparametric, for all biological replicates, considering the total number of colonies, mean±SEM, (*p=0.0155; **p=0.0092). BFU-E: burst forming unit erythroid; CFU-GM: colony forming unit granulocyte macrophage, CFU-M: colony forming unit macrophage; CFU-GEMM: colony forming unit granulocytes, erythrocytes, macrophages, megakaryocytes; **h**) Representative flow cytometric analysis of CD4^+^CD8^+^ T-cell potential of CD32^+^ (orange) and CD32^neg^ (blue) cells isolated at day 8 of WNTd hPSC-derived hematopoietic cultures. Gated on SSC/FSC/Live/CD45^+^CD56^neg^. H1 hESCs, *n*=3, independent; **i**) Bar plot showing the frequency of CD45^+^CD4^+^CD8^+^CD56^neg^ T-cells derived from CD32^+^ and CD32^neg^ fraction. One-tail paired Student’s t-test, nonparametric, for all biological replicates (H1 hESCs, *n*=3, independent), mean±SEM, **p=0.0073; **j**) Quantification of erythro-myeloid CFC potential of CD44^+^ (green) and CD44^neg^ (purple) populations isolated at day 8 of WNTd hPSC-derived hematopoietic cultures and cultured on OP9DLL1. One-tail paired Student’s t-test, nonparametric, for all biological replicates (H1 hESCs, *n*=3, independent) considering the total number of colonies, mean±SEM, **p=0.0098. BFU-E: burst forming unit erythroid; CFU-GM: colony forming unit granulocyte macrophage, CFU-M: colony forming unit macrophage; CFU-GEMM: colony forming unit granulocytes, erythrocytes, macrophages, megakaryocytes; **k**) Bar plot showing the frequency of CD45^+^ (yellow) and CD45^neg^ (grey) clones derived from CD32^+^ (orange) or CD44^+^ (green) cells isolated at day 8 of WNTd hPSC-derived hematopoietic cultures. Oneway ANOVA for all biological replicates (H1 hESCs, *n*=4), mean±SEM, (ns, not significant p=0.9855; ****p<0.0001; **p=0.0028; **l**) Table showing the clonal analysis of the day 8 WNTd hPSC-derived CD32^+^ (in orange, first columns) and CD44^+^ (in green, second column) cells cultured in the presence of pro-hematopoietic cytokines (HEC culture) as shown in Fig. 3k. Numbers indicate the number of wells seeded (first row), the number of wells showing clonal outgrowth (second row), the number of wells containing hematopoietic (CD45^+^, third row, in yellow) or non-hematopoietic (CD45^neg^, fourth row, in grey) cells relative to the total number of wells seeded. H1 hESCs, *n*=4, independent.

We therefore assessed the hematopoietic potential of CD34^+^CD43^neg^CD73^neg^CD184^neg^DLL4^neg^CD32^+/neg^ (referred to as CD32^+^ and CD32^neg^) cell populations (Fig. 3b, Extended Data Fig. 3a, c). The CD32^+^ cells generated erythro-myeloid clonogenic progenitors with significantly higher frequency than CD32^neg^ cells in both H1 and H9 hESC lines (Fig. 3g), similarly to what we showed in the human embryo (Fig. 2f). Given the residual hematopoietic potential observed in CD32^neg^ cells isolated from the AGM region and hPSC-derived hematopoietic cultures (Fig. 2f, 3g), we isolated CD32^neg^ cells from day 8 WNTd hematopoietic cultures and tested whether they could generate CD32^+^ progeny harboring hematopoietic potential. Indeed, the 2-day-long culture of CD32^neg^ cells gave rise to:::40% (39±1%) of CD32^+^ cells (Extended Data Fig. 3d) that, when isolated, generated CD45^+^ hematopoietic cells in HEC cultures (Extendend Data Fig. 3e). These data demonstrate that the CD32^neg^ cell population contains precursors of HECs that do not express CD32 yet. This evidence suggests that the residual hematopoietic output observed in the CD32^neg^ fraction is due to a further maturation of CD32^neg^ into CD32^+^ HECs that will then generate hematopoietic cells. We then assessed lymphoid potential by analyzing the defining lymphoid lineage for the WNTd hematopoietic program, *i*.*e*., T-cells^39^. Upon 24 days of coculture on OP9DLL4 stromal cells, only CD32^+^ cells could robustly generate CD4^+^CD8^+^ T-cells while the CD32^neg^ fraction did not display T-cell potential under the same conditions (Fig. 3h, i). Collectively, these results show that CD32 expression in CD34^+^CD43^neg^CD73^neg^CD184^neg^DLL4^neg^ cells demarcates a subpopulation endowed with robust multilineage intra-embryonic-like hematopoietic potential in hPSC cultures.

### CD32 is a specific HEC marker

We next compared the specificity of CD32 as hPSC-derived HEC marker against CD44, a surface marker often used to define HECs^21, 22^. In day 8 WNTd CD34^+^CD43^neg^ cells, CD44 is expressed by most DLL4^+^ cells (Extended Data Fig. 3f), in line with the reported CD44 expression in arterial endothelial cells^21, 25^, while it distinguishes two subpopulations of CD34^+^CD43^neg^CD184^neg^CD73^neg^DLL4^neg^ cells (referred to as CD44^+^ and CD44^neg^) (Extended Data Fig. 3g). We next assessed the hemogenic potential of CD44^+^ and CD44^neg^ subpopulations and observed that the CD44^+^ fraction was enriched for HECs as it generates significantly more clonogenic progenitors than CD44^neg^ cells (Fig. 3j), in accordance with recent reports^21, 22^. Given that both CD32 and CD44 expression enriches for cells with hemogenic potential, we analyzed the relationship between these two markers in hPSC differentiating cultures. Since CD32 identifies a minor subpopulation of CD44^+^ cells (Extended Data Fig. 3h) in day 8 WNTd CD34^+^CD43^neg^CD184^neg^CD73^neg^DLL4^neg^ cells, we hypothesized that CD32 might be a more specific marker for HECs than CD44. To compare the specificity of CD32 and CD44, we performed the single-cell HEC assay using CD32^+^ or CD44^+^ cells isolated at day 8 of WNTd hematopoietic cultures, as we have previously described^27^. This clonal analysis revealed that the CD32^+^ subfraction was highly enriched for HECs, as 87.0±4.0% of the cells that formed a clone (143/168) in the HEC assay generated exclusively CD45^+^ hematopoietic cells (Fig. 3k, l and Extended Data Fig. 3j). In stark contrast, the CD44^+^ fraction contained equal proportions of progenitors with either hematopoietic or non-hematopoietic potential (Fig. 3k, l). This single cell analysis indicates that CD32 is a reliable marker for hPSC-derived WNTd intra-embryonic HECs, as nearly all CD32^+^ cells harbor robust hematopoietic potential. In addition, the use of CD32 yields a significant improvement in enriching for HECs compared to CD44 (Fig. 3k, l), a marker often used to identify human HECs.

### CD32 identifies HECs in a specific Notch-independent state

In the attempt to further characterize hPSC-derived CD32^+^ cells, we next asked if these cells have transcriptional similarity to HECs found in the developing human embryo. We analyzed the transcriptomic profile of the CD32^+^ fraction sorted from day 8 WNTd hematopoietic cultures and, since HECs in the DA are found in close contact with non-hemogenic cells arterial cells^3^, we included CD34^+^CD43^neg^CD184^+^CD73^+^DLL4^+^ cells (referred to as DLL4^+^) as control sample^27^. While DLL4^+^ cells displayed a significant enrichment for genes whose expression is associated with arterial fate *in vivo* (*e.g.*, *CXCR4*, *DLL4*, *HEY1*, *HEY2*, *SOX17*, *GJA5*)^40^, CD32^+^ cells were enriched for the expression of genes associated with HECs and their hematopoietic progression *in vivo*, including *RUNX1*, *GFI1*, *MYCN* and *RAB27B* (Extended Data Fig. 4a and Supplementary Table 2)^40^. We then wondered whether *FCGR2B* expression could further refine the identification of HECs using available human CS14-15 AGM single cell RNA sequencing (scRNA-seq) data^40^. In this dataset, while *CDH5^+^RUNX1^+^PTPRC^neg^FCGR2B^neg^*HECs display an enrichment for genes associated with arterial endothelium, the expression of genes characteristic of hematopoietic commitment segregates within *CDH5^+^RUNX1^+^PTPRC^neg^FCGR2B^+^*expressing cells (Extended Data Fig. 4b). These data suggest that transcriptionally distinct states of HECs can be identified. To assess HEC heterogeneity and interrogate whether *FCGR2B* expression could define a specific state of HECs, we performed single-cell RNAseq (scRNAseq) of day 8 WNTd CD34^+^CD43^neg^CD184^neg^CD73^neg^ cells as they contain HECs as well as other endothelial progenitors and HEC precursors^27^. Unsupervised clustering revealed 22 transcriptionally distinct clusters which were mostly annotated to the major endothelial cell fates (Fig. 4a, Extended Data Fig. 4c and Supplementary Table 3a). This scRNAseq analysis confirmed that WNTd HECs showed transcriptional heterogeneity as a total of 6 clusters were enriched for cells differentially expressing *RUNX1* (Fig. 4a, Extended Data Fig. 4c, d and Supplementary Table 3a), including one with enriched expression of *FCGR2B* (cluster 11 in Fig. 4a, Extended Data Fig. 4e and Supplementary Table 3a).

**Figure 4.**
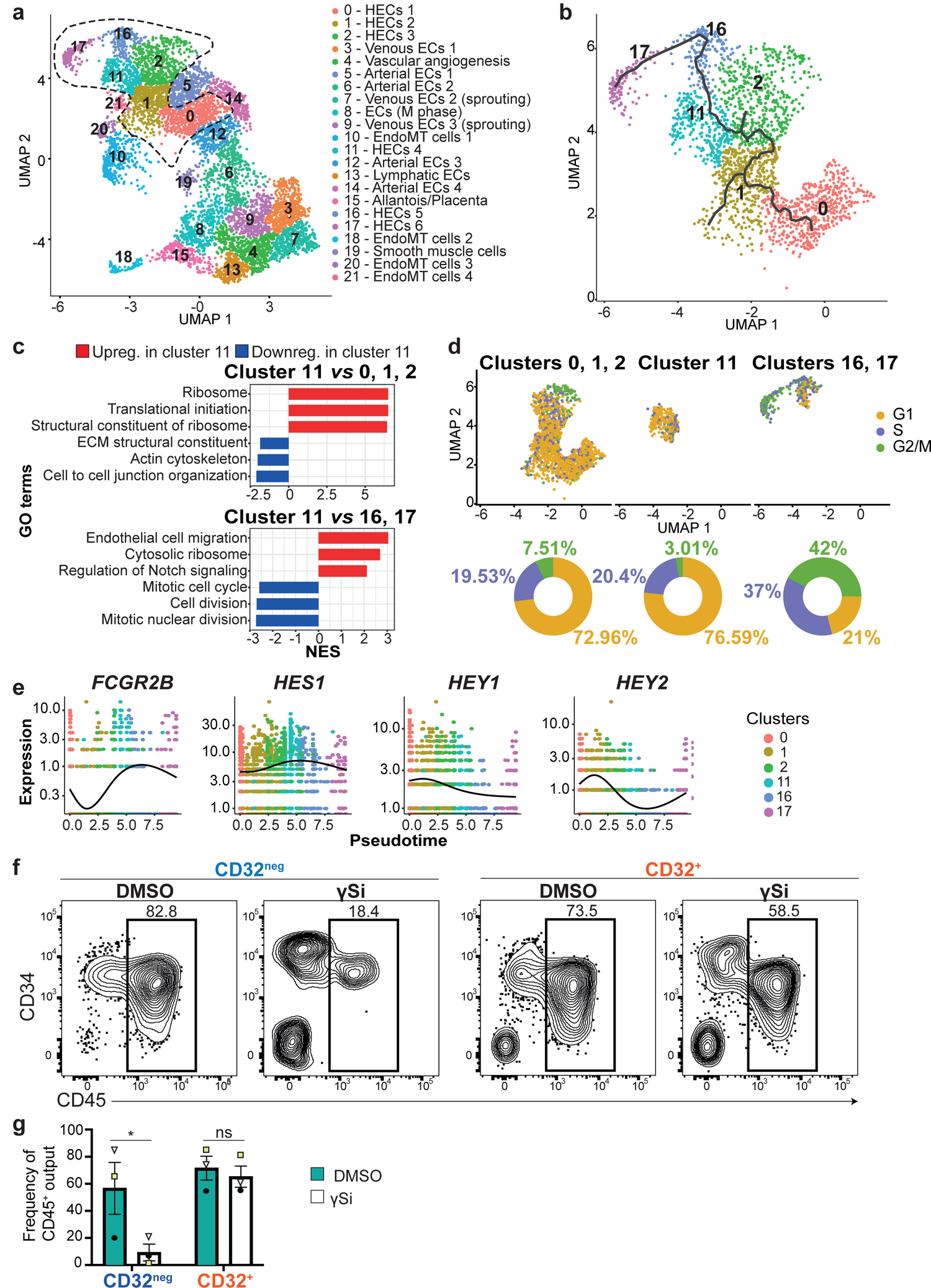
CD32 identifies a Notch-independent HEC state. **a**) UMAP of single-cell data of CD34^+^CD43^neg^CD184^neg^CD73^neg^ cells isolated at day 8 WNTd hPSC-derived hematopoietic cultures, H1 hESCs, *n*=1. Cells are clustered at resolution 0.6. *RUNX1* expressing clusters are highlighted by a black dashed line. HECs: hemogenic endothelial cells; ECs: endothelial cells; EndoMT: endothelial-to-mesenchyme transition; M phase: mitotic phase; **b**) UMAP visualization of the single-cell trajectory performed by Monocle3 on clusters 0, 1, 2, 11, 16, 17 that display *RUNX1* as a differential marker; **c**) Barplot showing significantly enriched Gene Ontology (GO) terms using GSEA (adjusted p-value<0.05) on fold change pre-ranked genes from the comparison between cluster 11 and clusters 0, 1, 2 (upper panel) or cluster 11 and clusters 16, 17 (bottom panel). Upregulated (upreg.) genes are shown in red, downregulated (downreg.) in blue. NES: normalized enrichment score, ECM: extracellular matrix: **d**) Upper panel: UMAP showing the cells in clusters 0, 1, 2, cluster 11, and cluster 16, 17 coloured according to the cell cycle phase. G1 phase is shown in yellow, S in purple, G2/M in green. Lower panel: donut charts showing the percentage of cells in G1, G2/M, and S phase in each of the clusters; **e**) Pseudotime kinetics of the expression variation of *FCGR2B* and the Notch targets genes *HES1, HEY1* and *HEY2* along the clusters with differential expression of *RUNX1* (clusters 0, 1,2, 11, 16, 17). Cells are coloured by the cluster identity. Lines denote relative average expression of each gene in pseudotime; **f**) Representative flow cytometric analysis of the hematopoietic CD45^+^ and endothelial CD34^+^ progeny derived after 4 days of HEC culture of CD32^+/neg^ cells isolated at day 8 of WNTd hPSC-derived hematopoietic cultures and treated with DMSO, as control, or γ-secretase inhibitor L-685,458 (γSi) to block Notch signaling. Gated on SSC/FSC/Live. *n*=3, independent; **g**) Bar plot showing the frequency of CD45^+^ cells derived from CD32^+^ and CD32^neg^ fraction treated with DMSO (in petroleum) or γ-secretase inhibitor L-685,458 (γSi, in white) to block Notch signaling. One-way ANOVA for all biological replicates (*n*=3), mean±SEM (*p=0.0456; ns, not significant, p=0.9623).

Pseudotime analysis by Monocle3 revealed that *FCGR2B^+^* HECs represent an intermediate state for the progression of *RUNX1*^+^*KCNK17*^+^*H19*^+^*FCGR2B*^neg^ HECs^41^ to *RUNX1*^+^ cells expressing hematopoietic genes such as *SPN* (despite being CD43^neg^) and *MYB* (Fig. 4b and Extended Data Fig. 4f, g). We next performed GSEA and ORA across this progression to dissect the unique features of *FCGR2B^+^* HECs and identify state-specific gene expression profiles proper of HECs with distinct characteristics. In particular, we observed that the gradual loss of endothelial identity of *H19*^+^ HECs begins with the downregulation of genes associated with extracellular organization, cell adhesion and cytoskeletal remodeling (Fig. 4c, Supplementary Table 3b-e). The progression to the *FCGR2B*^+^ HEC cluster is associated with an enrichment of the expression of several ribosomal protein genes, consistently with the role of RUNX1 in regulating ribosome biogenesis^42^, a key process for the emergence of blood cells (Fig. 4c, Supplementary Table 3b-e). Next, concomitantly to the upregulation of hematopoietic genes, HECs also display increased expression of genes associated with cell motility as well as cell cycle progression (Fig. 4c, Supplementary Table 3f-i). Indeed, using scRNA-seq data to infer the cell cycle state across HEC states, we observed that during the progression from *H19*^+^ to *FCGR2B*^+^ cluster (*i.e.,* from clusters 0, 1, 2 to cluster 11), cells are mostly in G1 phase, while cells in the *MYB*^+^ HEC clusters (*i.e*., clusters 16, 17) are mostly in S/G2/M phase (Fig. 4d).

Interestingly, the *MYB*^+^ HEC clusters (*i.e*., clusters 16, 17) negatively correlated with expression of genes associated to the activation of Notch signaling (Fig. 4c). Since Notch signaling is an essential driver of stage-specific intra-embryonic emergence of hematopoietic cells^12^, we further analyzed the expression trend according to pseudotime of the well characterized NOTCH target genes in the DA, *i.e., HES1*, *HEY1* and *HEY2* (Fig. 4e). The analysis revealed that *HES1* expression peaks in cells belonging to cluster 11, marked by *FCGR2B* differential expression, while *HEY1* and *HEY2* expression peaks in cells that precede *FCGR2B^+^* cells in this pseudotime (Fig. 4e). This suggests that *FCGR2B* expression might identify a Notch-independent state of HECs. To test this hypothesis, we added the chemical γ-secretase inhibitor L-685,458 (γSi) to the HEC culture of day 8 WNTd CD32^+^ as well as CD32^neg^ cells, as some of the latter will generate CD32^+^ cells (Extended Data Fig. 3e). While CD32^neg^ cells gave rise to hematopoietic progenitors in a Notch-dependent manner, the chemical inhibition of Notch signaling did not impair the generation of CD45^+^ hematopoietic progenitors from CD32^+^ cells (Fig. 4f, g). Altogether these results show that CD32 expression defines the temporal Notch-requirement within the HECs and that HECs activate a cell division program to give rise to hematopoietic progeny.

## Discussion

The precise identification of human HECs will enable the characterization of this transient population, unveil what regulates their transition to blood, thus allowing us to determine their identity, which is still subject to debate. In this study, using transcriptomic analysis of hemogenic populations sorted from human embryos, we identified *FCGR2B*, which encodes for a CD32 isoform, as a marker whose expression is upregulated in HECs. CD32 expression can be used in combination with other endothelial markers to precisely isolate HECs with robust hematopoietic potential from both the human embryo and hPSC differentiating cultures. In fact, CD32 expression is more specific for hPSC-derived HECs than CD44, another marker known to be expressed in HECs as well as in arterial cells^21, 22, 25^.

The use of a hPSC-based model has allowed us to capture and dissect in fine detail the HEC progression to the generation of blood cells. By providing new granularity to a rare process that *in vivo* occurs very rapidly, our study demonstrates that HECs can be found in different intermediate states, which display transcriptional profiles indicative of distinct cellular characteristics. The progression of HECs towards the hematopoietic fate begins with a gradual change in the expression of genes associated with the remodeling of the extracellular matrix, the loss of adhesion molecules and the re-organization of the cytoskeleton. This suggests that the transcriptional changes associated with the release of hematopoietic cells in the bloodstream occur before the actual morphological remodeling can be observed. In addition, this HEC progression culminates with the enhancement of ribosome biogenesis and translation in CD32^+^ HECs that appear to be irreversibly fated to hematopoiesis. Indeed, CD32^+^ HECs can generate hematopoietic progeny independently of Notch signaling, the major driver of this process. As such, our study defines the temporal requirement of Notch signaling to initiate the hematopoietic program in intra-embryonic HECs, with CD32 expression demarcating pre-*vs* post-Notch states during HEC progression to the blood fate.

While the initiation of the morphological remodeling and the concomitant expression of hematopoietic markers are often identified as the moment of the hematopoietic specification of HECs^43^, our data suggest that the commitment of HECs to the blood fate is temporally distinct from, and becomes Notch-independent before, the full execution of the hematopoietic program, which occurs at a different cell cycle state. In fact, while the fate decision coincides with a timely suppression of the cell cycle, the emergence of blood cells is associated with cell cycle re-entry. Given that the acquisition of an active cell cycle to generate lineage output is a hallmark of cell differentiation^44^, rather than developmental lineage transition^45, 46^, our findings support a model in which blood cells emerge via the differentiation of hematopoietic-restricted HECs rather than EHT.

In summary, this study demonstrates that expression of CD32 marks HECs fully committed to generate hematopoietic progeny and suggests that it could be used as a powerful tool to enrich hematopoietic precursors from a broad range of hPSC lines, including those for which current differentiation protocols into hematopoietic lineages are not optimal. Our findings will allow a deeper understanding of the specification of the HEC lineage, a central element of hematopoietic development, which will translate into optimized scaled-down, potentially cheaper, protocols to generate therapeutic blood products from hPSCs.

## Data availability

All gene expression analysis datasets are available in the Gene Expression Omnibus (GEO) under the accession number GSE223223.

## Code availability

Scripts used for data analysis and for the generation of all the figures in the paper are available at this link http://www.bioinfotiget.it/gitlab/custom/scarfo_hec2023.

## Acknowledgements

We thank Gordon Keller, Renato Ostuni, Luigi Naldini as well as C.M.S, M.T and A.D. lab members for inputs and critical reading of the manuscript; Marco Genua and Renato Ostuni for the help in library preparation, the Flow cytometry Resource, Advanced Cytometry Technical Applications Laboratory (FRACTAL), the Centro di Statistica per le Scienze Biomediche (CUSSB) at Ospedale San Raffaele and the Human Developmental Biology Resource. Sequencing was performed by the GenomEast platform (at the IGBMC Illkirch, France), a member of the “France Génomique” consortium (ANR-10-INBS-0009). M.T. was supported by INSERM and by grants from ANR (ANR-14-CE11-0008) and the Ligue Contre le Cancer Région Grand Est Bourgogne Franche Comté – CCIR Est. M.E.K. was awarded a fellowship from EURIdoc programme (H2020-MSCA-COFUND-2020). A.D. was supported by the Telethon Foundation (TIGET grant C4-2016/9 and TIGET grant G3b-2016/9) and San Raffaele Hospital (Seed Grant). R.S. conducted this study as partial fulfillment of an international Ph.D. in Molecular Medicine, Vita-Salute San Raffaele University. Human embryo and stem cell studies have been approved by the Ospedale San Raffaele Ethical Committee (TIGET-HPCT) and by the IRB Institutional of the French Institute of Medical Research and Health (Number 21-854).

## Author contributions

A.D. formulated the initial concept. R.S., M.E.K., J.N.F., C.M.S., M.T. and A.D. designed the experiments and analyzed the data; R.S., M.E.K., A.G., S.A.L, Z.L., S.C, E.D., S.A.L., M.T and A.D. performed the experiments; M.A.A., S.V., and I.M. performed bioinformatics analyses; R.S., C.M.S., M.T. and A.D. wrote the manuscript.

## Methods

### Human embryonic tissues

Human embryonic tissues employed for RNA sequencing, immunohistochemistry and immunofluorescence were obtained from voluntary abortions performed according to the guidelines and with the approval of the French National Ethics Committee. Written consent to the use of samples in research was obtained from patients. The study was approved by Ospedale San Raffaele Ethical Committee (TIGET-HPCT protocol) and by the Institutional Review Board of the French Institute of Medical Research and Health (Number 21-854). Human embryos were staged using anatomic criteria and the Carnegie classification. Samples were either used immediately as fresh tissues (*ex vivo* experiments and RNA sequencing analysis) or fixed in PBS supplemented with 4% paraformaldehyde (Sigma Aldrich), embedded in gelatin and stored at −80°C (immunohistochemistry and immunofluorescence). Human embryonic tissues (CS12-CS13) analyzed by RNA sequencing were incubated in medium containing 0.23% w/v collagenase Type I (Worthington Biochemical Corporation, NC9482366) for 30 minutes at 37°C and the single cell suspensions were filtered through a 70 µm cell strainer (BD Biosciences).

Human embryonic tissues (CS13) employed for *ex vivo* hematopoietic cultures were collected by human developmental biology resource (HDBR), Newcastle University, Newcastle, United Kingdom, with written informed consent and approval from the Newcastle and North Tyneside NHS Health Authority Joint Ethics Committee (08/H0906/21+5). The HDBR is regulated by the UK Human Tissue Authority (HTA; https://www.hta.gov.uk/) and operates in accordance with the relevant HTA Codes of Practice. The human embryonic tissues were dissociated for 50 minutes at 37°C with 10 mg/ml Collagenase/Dispase (Sigma Aldrich, 10269638001) in Phosphate Buffered Saline (PBS) with Ca^2+^ and Mg^2+^ (Sigma Aldrich, D8662), supplemented with 7% heat-inactivated fetal bovine serum (FBS, Hyclone, 12389802), 1% Penicillin-Streptomycin (Lonza, DE17-603E) and 10 µg/ml DNAse I (Calbiochem, 260913) and filtered through a 40 µm cell strainer (Falcon, 352235), similarly to what was previously described^47^.

### Immunohistochemistry and immunofluorescence

The techniques employed have been previously described^20^. Briefly, 5 μm sections were incubated first with primary antibodies overnight at 4°C, then for 1 hour at room temperature (RT) with biotinylated secondary antibodies and finally with fluorochrome-labeled (BioLegend) or peroxidase-labeled streptavidin (Beckman Coulter). Peroxidase activity was revealed with 0.025% 3,3-diaminobenzidine (Sigma Aldrich) in PBS containing 0.03% hydrogen peroxide. Low amounts of antigens (CD32, ACE) were revealed by Tyramide signal amplification (TSA) biotin or fluorescence amplification systems (Akoya, Biosciences). An isotype-matched negative control was performed for each immunostaining. When 3,3-diaminobenzidine was used on slides, they were counterstained with Gill’s hematoxylin (Sigma Aldrich), mounted in XAM neutral medium (BDH Laboratory Supplies), analyzed and imaged using an Optiphot 2 microscope (Nikon). Immunofluorescence-stained sections were cover-slipped in Prolong Gold Antifade Mountant with DAPI (Thermo Fisher Scientific) and analyzed with an Axio Imager M2 microscope coupled to a Hamamatsu’s camera Orca Flash 4v3 using the ApoTome.2 function (Zeiss) for optical sectioning.

The anti–human uncoupled primary antibodies used are listed: anti-human CD34 (Beckman, QBEnd/10), anti-human ACE (BB9, BD Biosciences, 557813), anti-human CD32 (Biolegend, 303202) and rabbit anti-human/mouse Runx1 (Abcam, ab92336). Secondary biotinylated antibodies were: goat anti-mouse IgG (Jackson Immuno Research, 115-066-072) and goat anti-rabbit IgG antibody (Jackson Immuno Research, 111-066-144). Double-immunofluorescence staining used the Dylight488 coupled streptavidin (Biolegend, 4052018) and the TSA Fluorescent Plus System.

### RNA sequencing data analysis

Human embryo sorted cells were collected in Eppendorf containing 6µl of PBS supplemented with 0.5µl of Protector RNAse inhibitor (Roche, 3335399001) and conserved in −80°C. Full length cDNA was generated using Clontech SMART-Seq v4 Ultra Low Input RNA kit for Sequencing (Takara Bio Europe, 634891) according to manufacturer’s instructions with 15 cycles of PCR for coding DMA (cDNA) amplification by Seq-Amp polymerase. 600 pg of pre-amplified cDNA were then used as input for Tn5 transposon tagmentation by the Nextera XT DNA Library Preparation Kit (Illumina, FC-131-1096) followed by 12 cycles of library amplification. After purification with Agencourt AMPure XP beads (Beckman-Coulter, A63882), the size and concentration of libraries were assessed by capillary electrophoresis. Libraries were then sequenced with the Illumina HiSeq 4000 sequencing platform in the single-end mode and with a read length of 50 base pairs (bp). For day 8 WNTd hPSC-derived hematopoietic cultures, total RNA from sorted CD34^+^CD32^+^CD43^neg^CD184^neg^CD73^neg^DLL4^neg^ and CD34^+^CD184^+^CD73^+^DLL4^+^CD43^neg^ was purified using the ReliaPrep RNA Cell Miniprep System and RNA-Seq libraries were generated using the Smart-seq2 method (Picelli et al., 2014). 1 ng of RNA were retrotranscribed, cDNA was PCR-amplified (15 cycles) and purified with AMPure XP beads. After purification, the concentration was determined using Qubit 3.0 and size distribution was assessed using Agilent 4200 TapeStation system. Then, the tagmentation reaction was performed starting from 0.5 ng of cDNA for 30 minutes at 55°C and the enrichment PCR was carried out using 12 cycles. Libraries were then purified with AMPure XP beads, quantified using Qubit 3.0, assessed for fragment size distribution on an Agilent 4200 TapeStation system. Sequencing was performed on an Illumina NovaSeq6000 (single-end, 100 bp read length) following manufacturer’s instruction.

For both RNA sequencing datasets, raw reads quality control was accomplished using the FastQC tool (http://www.bioinformatics.babraham.ac.uk/projects/fastqc) and read trimming was performed using the Trim Galore software (https://doi.org/10.5281/zenodo.5127899) to remove residual adapters and low-quality sequences. Trimmed reads were aligned against the human reference genome (GRCh38) using STAR^48^ with standard parameters. Uniquely mapped reads were then assigned to genes using the featureCounts tool from the Subread package^49^, considering the GENCODE primary assembly v.34 gene transfer file (GTF) as reference annotation for the genomic features. Gene count matrices were then processed by using the R/Bioconductor differential gene expression analysis packages DESeq2^50^ applying the standard workflow.

For the human embryos’ dataset, a paired analysis was set up modeling gene counts using the following design formula: ∼donor + condition. Gene p-values were corrected for multiple testing using FDR. Genes with adjusted p-values < 0.05 were considered differentially expressed.

Over representation analysis (ORA) and a gene-set enrichment analysis (GSEA) were then computed considering the Gene Ontology (GO) Biological Process (BP) terms from the C5 collection of the Molecular Signatures Database (MSigDB version 7.2) using the R/Bioconductor package clusterProfiler^51^ (v 3.8.1, http://bioconductor.org/packages/release/bioc/html/clusterProfiler.html). ORA was applied to the significantly differentially expressed genes, while GSEA was performed by pre-ranking genes according to fold change values. P-values were corrected for multiple testing using FDR and enriched terms with an adjusted p-value less than 0.05 were considered statistically significant. Volcano plots were generated using the R package ggplot2 (https://ggplot2.tidyverse.org) and have been used to display RNA-seq results plotting the statistical significance (adjusted P-value) *vs* the magnitude of change (fold change). Heatmaps were generated using the R package pheatmap (https://CRAN.R-project.org/package=pheatmap). Surface genes were extracted using the surfaceome database (Bausch-Fluck *et al*, 2018) (http://wlab.ethz.ch/surfaceome/)^52^.

### Single-cell RNA-seq analysis

Cells from WNTd hPSC-derived day-8 hematopoietic culture condition were methanol-fixed as previously described^53^. Libraries were prepared following the manufacturer’s instructions using the Chromium platform (10x Genomics, Pleasanton, CA) with the 3’ gene expression (3’ GEX) V3 kit, using an input of ∼10,000 cells. Briefly, Gel-Bead in Emulsions (GEMs) were generated on the sample chip in the Chromium controller. Barcoded cDNA was extracted from the GEMs by Post-GEM RT-cleanup and amplified for 12 cycles. Amplified cDNA was fragmented and subjected to end-repair, poly A-tailing, adapter ligation, and 10X-specific sample indexing following the manufacturer’s protocol. cDNA libraries were quantified using Bioanalyzer (Agilent) and QuBit (Thermofisher) analysis and were sequenced in paired end mode on a NovaSeq instrument (Illumina, San Diego, CA) targeting a depth of 50,000-100,000 reads per cell. Sequencing reads were processed into gene count matrix by Cell Ranger (https://support.10xgenomics.com/single-cell-gene-expression/software/pipelines/latest/what-is-cell-ranger, v4.0.0) from the Chromium Single Cell Software Suite by 10x Genomics. In detail, fastq files were generated using the Cell Ranger ‘*mkfastq’* command with default parameters. Gene counts for each cell were quantified with the Cell Ranger *‘count’* command with default parameters. The human genome (GRCh38.p13) was used as the reference. The resultant gene expression matrix was imported into the R statistical environment (version 4.0.3) for further analyses. Cell filtering, data normalization, and clustering were carried out using the R package Seurat (Stuart et al., 2019) v3.2.2. For each cell, the percentage of mitochondrial genes, number of total genes expressed and cell cycle scores (S and G1 phase) were calculated. Cells with a ratio of mitochondrial vs. endogenous gene expression > 0.2 were excluded as putative dying cells. Cells expressing <200 or >6,000 total genes were also discarded as putative poorly informative cells and multiplets, respectively. Cell cycle scores were calculated using the ‘CellCycleScoring’ function that assigns to each cell a score based on the expression of the S and G2/M phase markers and stores the S and G2/M scores in the metadata along with the predicted classification of the cell cycle state of each cell. Counts were normalized using Seurat function ‘*NormalizeData’* with default parameters. Expression data were than scaled using the ‘*ScaleData’* function, regressing on the number of unique molecular identifier, the percentage of mitochondrial gene expression, and the difference between S and G2M scores. By using the most variable genes, dimensionality reduction was then performed with principal component analysis (PCA) by calculating 100 PCs and selecting the top 55 PCs. Uniform Manifold Approximation and Projection (UMAP) dimensionality reduction (McInnes et al., 2018) was performed on the calculated principal components to obtain a 2D representation for data visualization. Cell clusters were identified using the Louvain algorithm at resolution r = 0.6, implemented by the ‘*FindCluster’* function of Seurat. To find the differentially expressed (marker) genes from each cluster, the ‘*FindAllMarkers’* function (iteratively comparing one cluster against all the others) from the Seurat package was used with the following parameters: adjusted P values <0.05, average log FC >0.25, and percentage of cells with expression > 0.1. A comprehensive manual annotation of the cell types was performed using the previously obtained markers list. Differentially expressed genes between cells of cluster 11 against cells of clusters 0,1, and 2 and clusters 16 and 17 were determined by the ‘FindMarkers’ function using the following paramaters adjusted P values <0.05, |average log FC| > 0, and percentage of cells with expression > 0. GSEA was then performed considering Gene Ontology (GO) Biological Process (BP) terms from the C5 collection of the Molecular Signatures Database (MSigDB version 7.2) using the R/Bioconductor package clusterProfiler^51^ (v 3.8.1, http://bioconductor.org/packages/release/bioc/html/clusterProfiler.html). ORA was computed on the significantly differentially expressed genes considering Gene Ontology (GO) Biological Process (BP) terms from the C5 collection of the Molecular Signatures Database (MSigDB version 7.2) and the Reactome Pathways Database using the R/Bioconductor package ^51^ (v 3.8.1). P-values were corrected for multiple testing using FDR and enriched terms with an adjusted p-value less than 0.05 were considered statistically significant. Barplot was constructed using the R package ggplot2 (https://ggplot2.tidyverse.org).

Single-cell RNA-seq samples from the public dataset GSE162950 were retrieved and processed as described in ^40^. The ‘*DotPlot’* function from the Seurat R package was used to construct a scorecard highlighting the expression pattern of selected cells having RUNX1+ CDH5+ FCGR2B+ or RUNX1 CDH5+ FCGR2B-expression patterns.

Pseudotime trajectory was constructed using Monocle3 (version 0.2.3) (https://cole-trapnell-lab.github.io/monocle3/)^54, 55^. Expression and feature data were extracted from the Seurat object and a Monocle3 ‘*cell_data_set’* object was constructed. The processed data was normalized followed by Principal Component Analysis (PCA) analysis using the Monocle3 function ‘*preprocess_cds’*. Dimensionality reduction was performed using the ‘*reduceDimension’* function. Trajectory graph learning and pseudo-time measurement through reversed graph embedding were performed with ‘*learn_graph’* function. Cells were ordered along the trajectory using the ‘*orderCells’* method with default parameters. The ‘*plot_cells’* function was used to generate the trajectory plots.

### Human pluripotent stem cells maintenance and differentiation

The use of human embryonic stem cells (hESCs) was approved by the Ospedale San Raffaele Ethical Committee, included in the TIGET-HPCT protocol. The already established H1^56^ and *RUNX1C*-EGFP H9^30^ hESCs lines were grown on irradiated mouse embryonic fibroblast (MEF) feeders in hES medium defined as DMEM/F12 medium (Corning, L022046-10092CVR) supplemented with 25% of KnockOut™ Serum Replacement (Thermo Fisher Scientific, 10828028), 1% Penicillin-Streptomycin (Lonza, DE17-603E), 2 mM L-Glutamine (Lonza, BE17-605E), 0.1% β-Mercaptoethanol (Sigma Aldrich, M3148), 0.7% of MEM Non-Essential Amino Acids Solution (Thermo Fisher Scientific, 11140035). 1 µg/ml Ciprofloxacin HCl (Sigma Aldrich, PHR1044-1G) and 20 ng/ml human recombinant basic fibroblast growth factor (bFGF, R&D, 233-FB-500/CF) were added to hES medium right before usage. Cells were maintained and expanded at 37°C, 21% O2, 5% CO2.

For differentiation, hPSC were cultured on Matrigel-coated plasticware (Corning Life Sciences, 356230) for 24 hours, followed by embryoid body (EB) generation. EB aggregates were resuspended in SFD media defined as 75% IMDM (Corning, 15343531), 25% Ham’s F12 (Corning, 10-080-CVR), 0.005% BSA – Fraction V, B27 supplement (Thermo Fisher Scientific Cat # 12587010), N2 supplement (Thermo Fisher Scientific Cat # 17502048), 1% Penicillin-Streptomycin, 1 µg/ml Ciprofloxacin HCl. The differentiation media was supplemented as previously described^27, 57^. Briefly, the first day of differentiation SFD medium was supplemented with 2 mM L-glutamine, 1 mM ascorbic acid (Sigma Aldrich, A4544), 400 µM 1-Thioglycerol solution (Sigma-Aldrich, M6145), 150 μg/ml transferrin (R&D, 2914-HT), and 10 ng/ml BMP4 (R&D, 314-BP-MTO). 24 hours later, 5 ng/ml bFGF (R&D, 233-FB-500/CF) was added. At the second day of differentiation, 3 μM CHIR99021 (Cayman Chemical Company, CT99201) were added, as indicated. On the third day, embryoid bodies (EBs) were changed to StemPro-34 media (Thermo Fisher Scientific, 10639011) supplemented with Penicillin-Streptomycin, L-glutamine, ascorbic acid, 1-Thioglycerol and transferrin, as above, with additional 5 ng/ml bFGF and 15 ng/ml VEGF (R&D, MAB3572). On day 6, 10 ng/ml IL-6 (130-093-934), 25 ng/ml IGF-1 (130-093-887), 5 ng/ml IL-11 (130-103-439), 50 ng/ml SCF (130-096-696), 2 U/ml EPO (Peprotech, 100-64) were added. To assess the emergence of CD32^+^ cells from CD32^neg^, CD32^neg^ cells were isolated at day 8 of WNTd hematopoietic cultures and cultured in day 6 media for 48 hours. All cytokines were purchased from Miltenyi Biotec, unless indicated differently. All differentiation cultures were maintained at 37°C. All embryoid bodies and mesodermal aggregates were cultured in a 5% CO_2_, 5% O_2_, 90% N_2_.

### T-cell differentiation

To test the T-cell potential, candidate cells isolated by FAC-sorting as indicated were seeded on OP9DLL4- or OP9DLL1-coated 24-well plates. OP9 cells expressing murine DLL1 or human DLL4 were a kind gift from Juan-Carlos Zúñiga-Pflücker and described previously^58, 59^. The cells were cultured in Alpha MEM (Thermo Fisher, 12000063) supplemented with 2.2 g/L sodium bicarbonate (Corning, 61-065-RO), 20% FBS (HyClone), 1% Penicillin-Streptomycin, 2mM glutamine (Thermo Fisher Scientific) and 400 µM 1-Thioglycerol solution supplemented with lineage specific cytokines. Cells were supplemented 5 ng/ml IL7, 5ng/ml FLT3L and, for the first 5 days of differentiation, 50 ng/ml SCF. The cells were split every 4-5 days by vigorous pipetting and passaging through a 40μm cell strainer and plated on freshly seeded OP9DLL4. T-lymphoid output was assayed by FACS analysis after 21-24 days of differentiation. Cells were analyzed using a LSR Canto II flow cytometer (BD).

### Colony forming cell (CFC) generation assay

The CFC generation assay was performed as previously described^60^. Briefly, sorted cells were cultured on irradiated OP9DLL1 monolayers in Alpha-MEM (Alpha-MEM, ThermoFisher, 12000063) supplemented with 20% FBS (HyClone), 1% Penicillin-Streptomycin 2 mM L-glutamine, 30 ng/ml TPO (Miltenyi Biotec, 130-095-747), 10 ng/ml BMP4, 50 ng/ml, 25 ng/ml IGF1, 10 ng/ml IL11, 10 ng/ml FLT3L, 4 U/ml EPO. After five days, cells were collected using 0.25% Trypsin-EDTA (Thermo Fisher Scientific, 25-200-056) for 3’ at 37°C. Cells were then filtered through a 40 μM filter and seeded on methylcellulose medium (Stemcell Technologies, H4034). Cells were seeded on methylcellulose supplemented with 150 ug/ml transferrin, 50 ng/ml TPO, 10 ng/ml VEGF, 10 ng/ml IL6, 50 ng/ml IGF1, 5 ng/ml IL11, 4 U/ml EPO. Colonies’ number and morphology was evaluated after fifteen days by light microscopy.

### HEC culture

CD34^+^CD43^neg^CD184^neg^CD73^neg^DLL4^neg^CD32^+/neg^ were isolated at day 8 of WNTd hematopoietic culture and reaggregated overnight at 3×10^5^ cells/ml as previously described^27^. The cells were seeded in Stempro media, supplemented with 1% glutamine, 50 μg/ml ascorbic acid, 150 μg/ml transferrin, 400 μM 1-Thioglycerol solution, 30 ng/ml TPO, 10 ng/ml VEGF, 5 ng/ml bFGF, 30 ng/ml IL3, SCF, 50 ng/ml IGF1, 10 ng/ml IL6, 5 ng/ml IL11, 4 U/ml EPO. Aggregates were then transferred onto thin-layer Matrigel-coated plasticware where they were cultured for an additional 1-7 days in the same media.

10 μM γ-secretase inhibitor (γSi) L-685,458 (Tocris, 2627) or equal volume of DMSO (D4540, Sigma Aldrich), as control, were added to the HEC culture test the effect of Notch signaling inhibition.

Single CD34^+^CD43^neg^CD184^neg^CD73^neg^DLL4^neg^CD32^+^/CD44^+^ cells were FAC-sorted directly onto a Matrigel-coated well of 96-well plate at day 8 of WNTd hematopoietic cultures. Cells were cultured as above. Hematopoietic and non-hematopoietic clones were evaluated by light microscopy and FACS analysis after 10-14 days of culture.

### Cell staining, flow cytometry and cell sorting

Samples for FACS analysis or cell sorting were incubated with antibody mixes for 15-30’ at 4°C. The antibodies employed are listed in the table below. Dead cells were excluded using 7AAD during staining. Cells were sorted with FACSAria II (BD). Sorting gates were set using appropriate fluorescence minus one (FMO) and single staining controls. FACS-analysis were performed either at FACS Canto (BD Biosciences) or Cytoflex S (Beckman Coulter).

### Antibodies list

| Anti-human antibodies |  |  |  |  |
| --- | --- | --- | --- | --- |
| ACE (BB9) | - | BD | 557813 | 2:100 |
| CD184 | BV421 | BD | 562448 | 1:100 |
| CD32 | APC Cyanine 7 | BD | 303229 | 2:100 |
| CD32 | PE | BD | 303206 | 2:100 |
| CD34 | APC | Beckman Coulter | IM2472 | 2:100 |
| CD34 | PE Cyanine 7 | Biolegend | 343616 | 1:400 |
| CD34 | PE Cyanine 7 | eBioscience | 25034942 | 1:400 |
| CD4 | PE Cy7 | BD | 560649 | 2:100 |
| CD43 | APC | BD | 560198 | 2:100 |
| CD43 | FITC | BD | 555475 | 10:100 |
| CD44 | PE | Miltenyi | 130-113-342 | 2:100 |
| CD45 | APC Cyanine 7 | Biolegend | 368516 | 1:100 |
| CD45 | FITC | Beckman Coulter | A07782 | 2:100 |
| CD45 | BV421 | BioLegend | 304032 | 2:100 |
| CD5 | PE | BioLegend | 300607 | 5:100 |
| CD56 | BV421 | BD | 740076 | 3:100 |
| CD7 | APC | BD | 561604 | 2:100 |
| CD73 | BV421 | BD | 562430 | 3:100 |
| CD8 | PE | Biolegend | 31051 | 2:100 |
| DLL4 | APC | Biolegend | 346508 | 1:100 |
| DLL4 | BV421 | BD | 744840 | 1:100 |
| anti-IgG1 | PE | Southern Biotechnology | 1070-05 | 1:1000 |
| Fcr Blocking reagent human | - | Miltenyi | 130-059-901 | 20:100 |
|  | 7AAD | BD | 559925 | 1:100 |

## Extended Data Figure legends

**Extended Data Figure 1.**
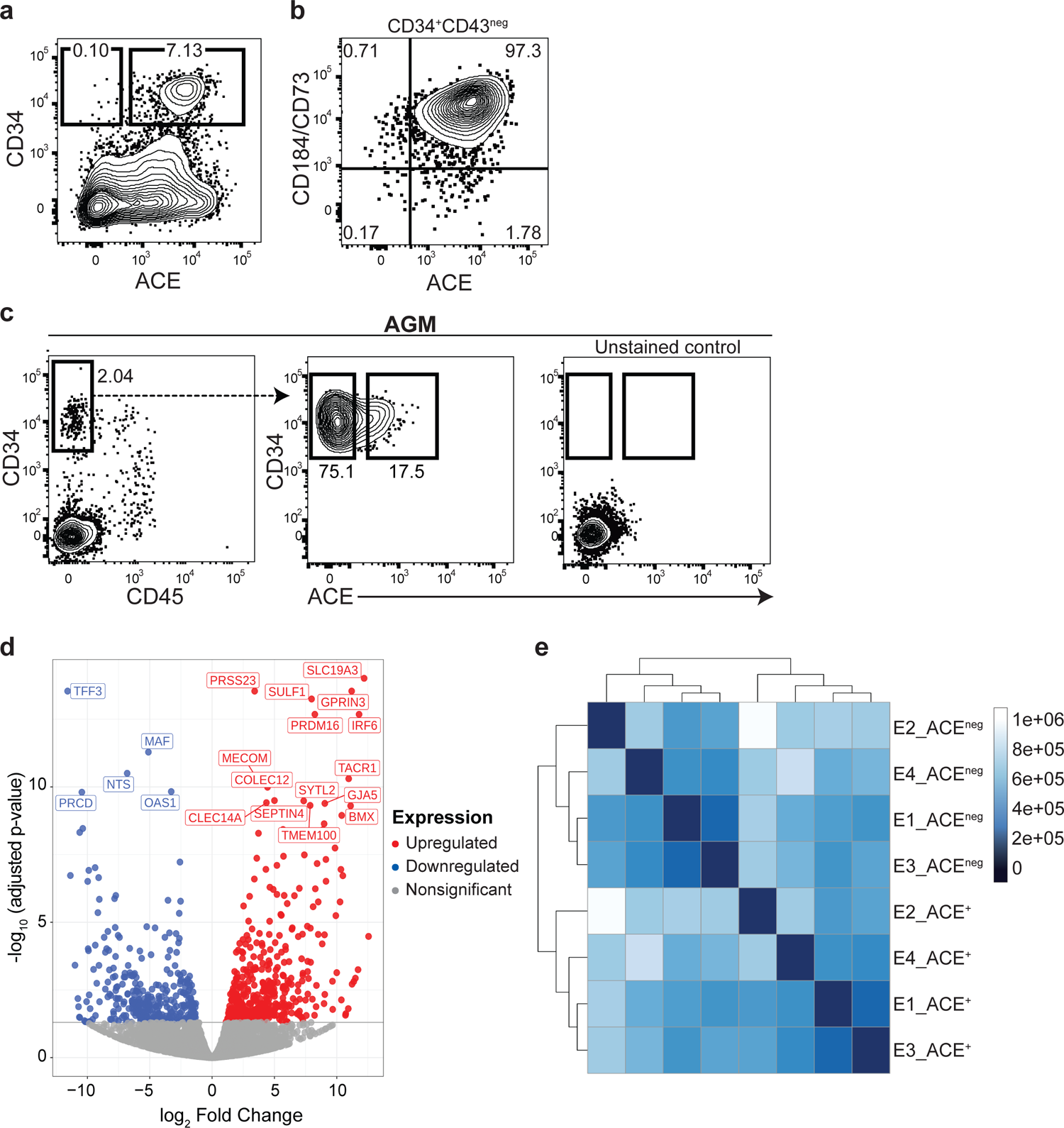
ACE is expressed within CD34^+^ cells during intra-embryonic hematopoiesis. **a**) Representative flow cytometric analysis showing ACE and CD34 expression in day 8 WNTd hPSC-derived hematopoietic cultures. Gated on SSC/FSC/Live. H1 hESCs, *n*=4, independent; **b**) Representative flow cytometric analysis showing CD184/CD73 and ACE expression in CD34^+^CD43^neg^ cells at day 8 of WNTd hPSC-derived hematopoietic cultures. Anti-CD184 and anti-CD73 antibodies are in the same color. Gated on SSC/FSC/Live/CD34^+^CD43^neg^. H1 hESCs, *n*=4, independent; **c**) Representative flow cytometric analysis showing the gating strategy to isolate ACE^+^ and ACE^neg^ cells from the AGM of *n*=4 CS12-CS13 human embryos. Left panel: CD45 and CD34 expression, gated on SSC/FSC/Live. Middle panel: ACE and CD34 expression, gated on SSC/FSC/Live/CD34^+^CD45^neg^. Right panel: unstained control, gated on SSC/FSC/Live; **d**) Volcano plot showing the differentially expressed genes in ACE^+^ and ACE^neg^ cells isolated from the AGM of four CS12-CS13 human embryos. The top 20 differentially expressed genes are highlighted, upregulated in red, downregulated in blue, nonsignificant in grey; **e**) DESeq2 heatmap distance analysis of four CS12-CS13 human embryos used for RNA sequencing. Samples were clustered using unsupervised hierarchical clustering by K-Means.

**Extended Data Figure 2.**
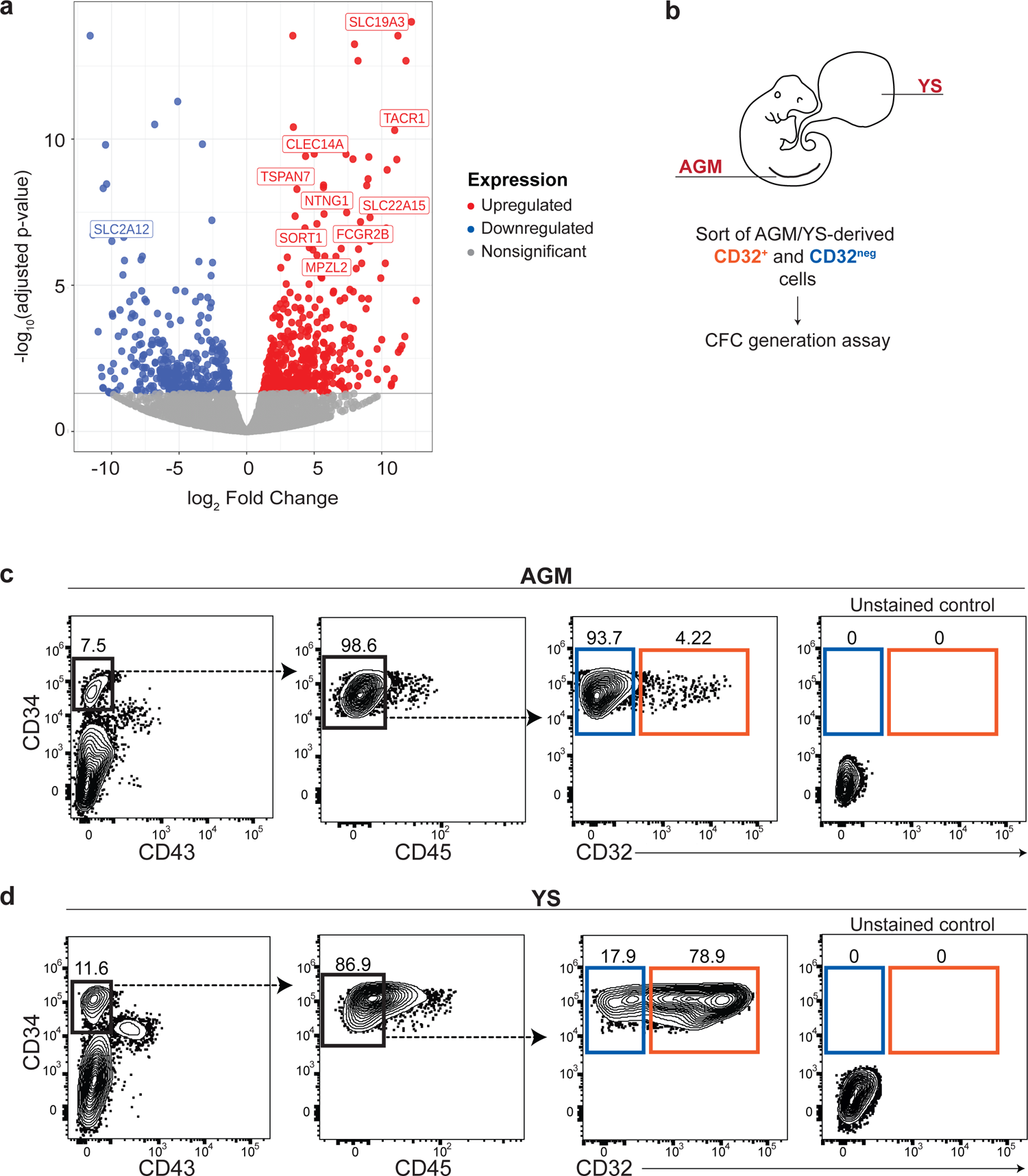
Identification of CD32 in the AGM and the YS of human embryos. **a**) Volcano plot showing the differentially expressed cell surface genes in ACE^+^ vs ACE^neg^ cells isolated from the AGM of four CS12-CS13 human embryos. The top 10 differentially expressed cell surface genes are highlighted, upregulated in red, downregulated in blue, nonsignificant in grey; **b**) Experimental layout: CD34^+^CD43^neg^CD45^neg^CD32^+/neg^ (referred to as CD32^+^ and CD32^neg^) cells were FAC-sorted from the AGM and YS of two CS13 human embryos. Isolated cells were tested for their hematopoietic potential via CFC generation assay; **c**) Representative flow cytometric analysis showing the gating strategy to isolate CD32^+^ (orange) and CD32^neg^ (blue) cells from the AGM of CS13 human embryo. From the left, first panel: gated on SSC/FSC/Live. Second panel: gated on SSC/FSC/Live/CD34^+^CD43^neg^. Third panel: gated on SSC/FSC/Live/CD34^+^CD43^neg^CD45^neg^. Fourth panel: unstained control, gated on SSC/FSC/Live. n=2, independent; **d**) Representative flow cytometric analysis showing the gating strategy to isolate CD32^+^ (orange) and CD32^neg^ (blue) cells from the YS of CS13 human embryo. n=2, independent. From the left, first panel: gated on SSC/FSC/Live. Second panel: gated on SSC/FSC/Live/CD34^+^CD43^neg^. Third panel: gated on SSC/FSC/Live/CD34^+^CD43^neg^CD45^neg^. Fourth panel: unstained control, gated on SSC/FSC/Live.

**Extended Data Figure 3.**
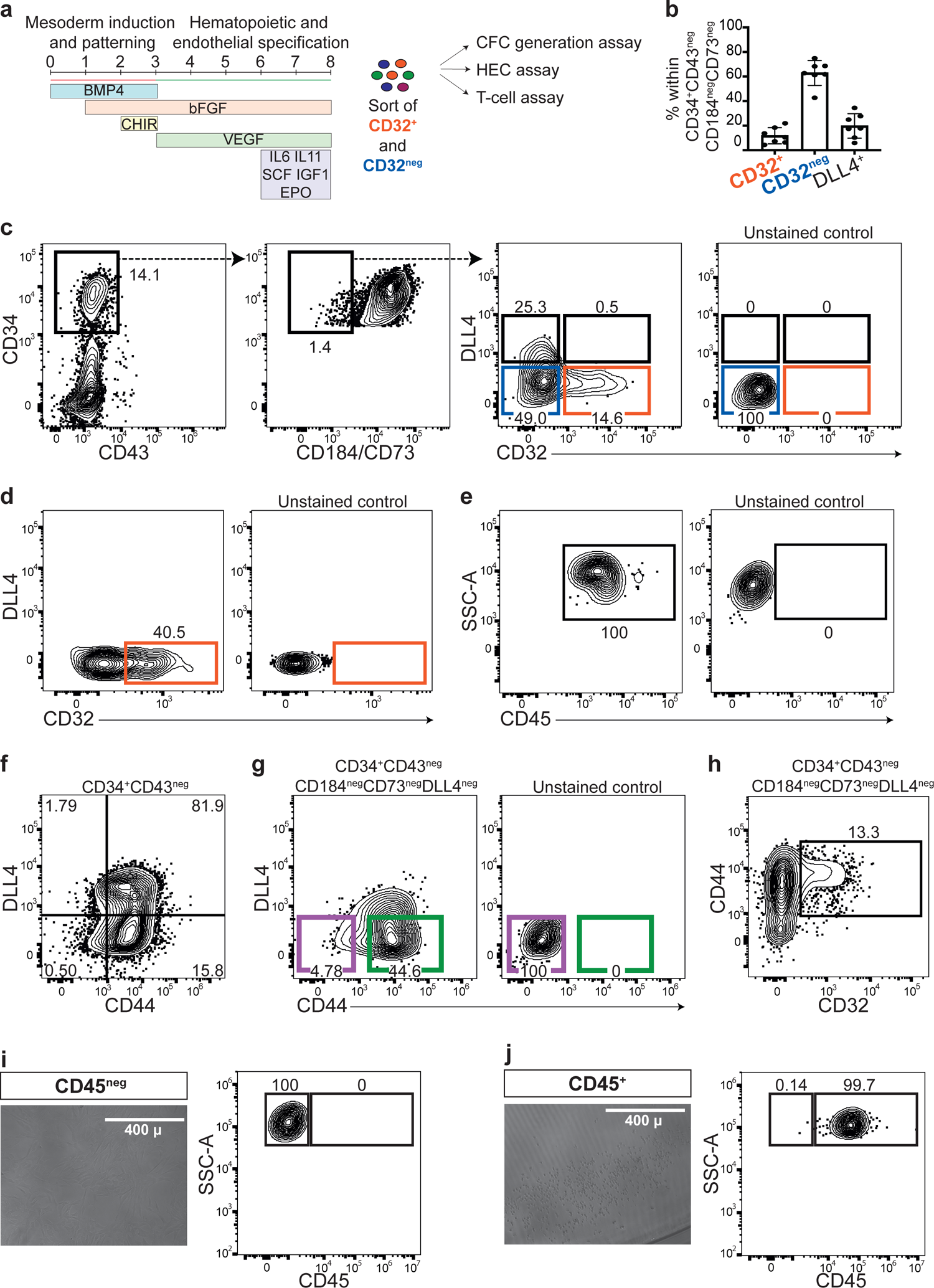
Characterization of the CD32^+^ and the CD44^+^ cell populations in WNTd hPSC hematopoietic cultures. **a**) Experimental layout showing the timeline of WNTd hPSC-derived hematopoietic cultures obtained by adding the WNT agonist CHIR 99021. CD34^+^CD43^neg^CD184^neg^CD73^neg^DLL4^neg^CD32^+/neg^ (referred to as CD32^+^, CD32^neg^) cells were isolated at day 8 and further cultured to assay the CFC generation, T-lymphoid potential or the generation of hematopoietic progeny through HEC culture; **b**) Bar plot showing the frequency of DLL4^neg^CD32^+/neg^ cells and DLL4^+^ within day 8 of WNTd hPSC-derived CD34^+^CD43^neg^CD184^neg^CD73^neg^ cells. Mean±SEM. H1 hESCs *n*=7, independent; **c**) Representative flow cytometric analysis showing the gating strategy to isolate CD32^+^ (orange) and CD32^neg^ (blue) cells from day 8 of WNTd hPSC-derived hematopoietic cultures. DLL4^+^ cells are highlighted in black. From the left, first panel: gated on SSC/FSC/Live. Second panel: gated on SSC/FSC/Live/CD34^+^CD43^neg^. Third panel: gated on SSC/FSC/Live/ CD34^+^CD43^neg^CD184^neg^CD73^neg^. Fourth panel: unstained control, gated on SSC/FSC/Live. *n*=7, independent; **d**) Representative flow cytometric analysis of the generation of CD32^+^ cells from CD32^neg^ cells isolated at day 8 of WNTd hPSC-derived hematopoietic cultures and cultured for 2 extra days using the same culture conditions. Gated on SSC/FSC/Live/CD34^+^CD43^neg^CD184^neg^CD73^neg^. H9 hESCs, *n*=3, independent; **e**) Representative flow cytometric analysis of the hematopoietic CD45^+^ progeny derived from CD32^+^ cells isolated 2 days after the sorting of the CD32^neg^ fraction at day 8 of WNTd hPSC-derived hematopoietic cultures as shown in d). Gated on SSC/FSC/Live. H9 hESCs, *n*=3, independent; **f**) Representative flow cytometric analysis of CD44 and DLL4 expression in day 8 WNTd hPSC-derived CD34^+^CD43^neg^ cells. Gated on SSC/FSC/Live/CD34^+^CD43^neg^. H1 hESCs, *n*=4, independent; **g**) Representative flow cytometric analysis of CD44 and DLL4 expression in day 8 WNTd hPSC-derived CD34^+^CD43^neg^CD184^neg^CD73^neg^ cells. Two populations are highlighted within DLL4neg fraction: CD44^+^ in green and CD44^neg^ in purple. Left panel: gated on SSC/FSC/Live/ CD34^+^CD43^neg^CD184^neg^CD73^neg^.Right panel, unstained control, gated on SSC/FSC/Live. H1 hESCs, *n*=4, independent; **h**) Representative flow cytometric analysis of CD32 and CD44 expression in day 8 WNTd hPSC-derived hematopoietic cultures. Gated on SSC/FSC/Live/CD34^+^CD43^neg^CD184^neg^CD73^neg^DLL4^neg^. *n*=3, independent; **i**) and **j**) Photo-micrograph (left panel) and representative CD45 flow cytometric analysis of a clone composed by adherent non-hematopoietic cells (**i**) or round hematopoietic cells (**j**) derived from single CD32^+^ cell isolated at day 8 of WNTd hPSC-derived hematopoietic culture. Gated on SSC/FSC/Live. H1 hESCs, *n*=3, independent.

**Extended Data Figure 4.**
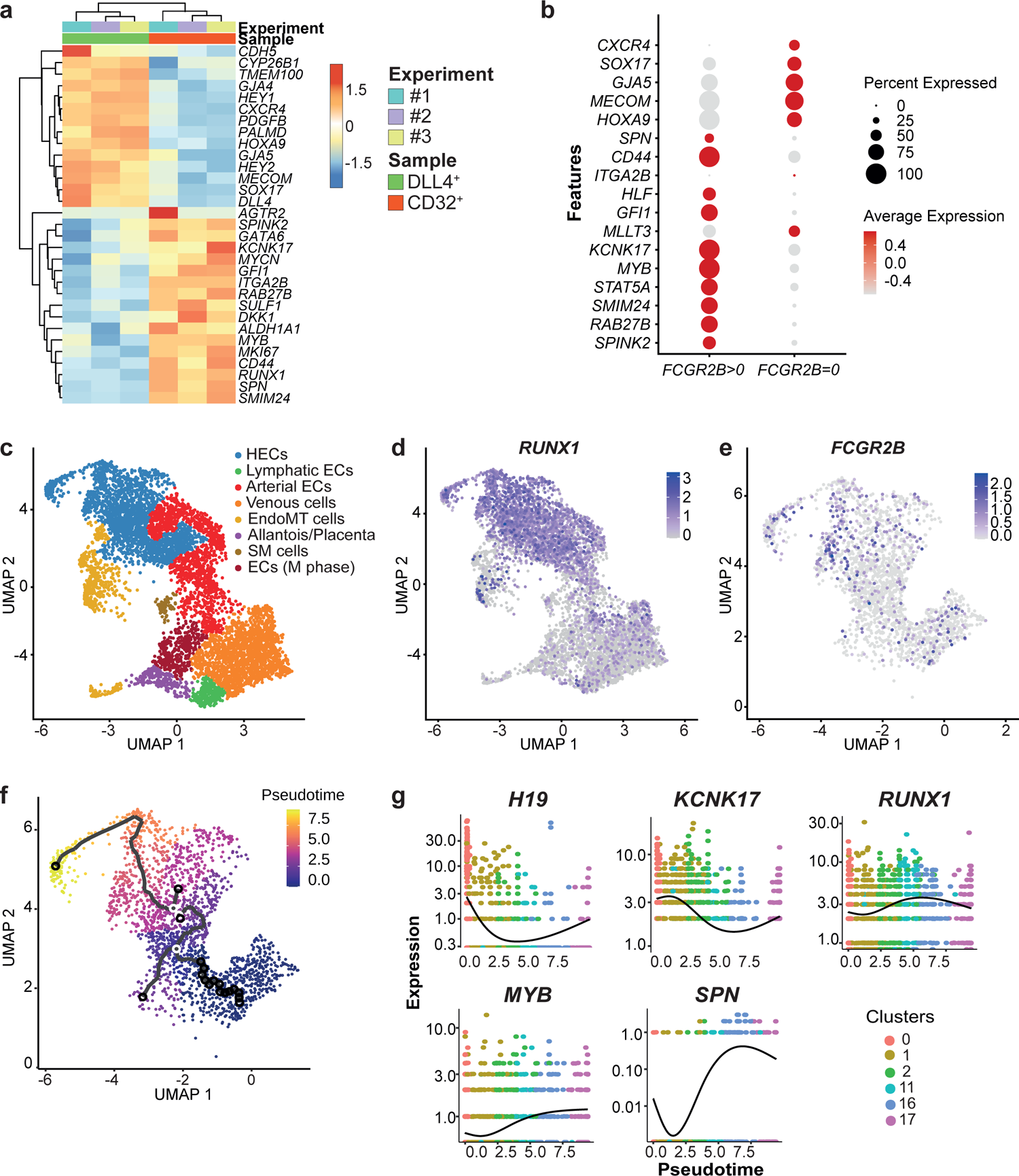
Transcriptomic analysis identifies the heterogeneity of hPSC-derived HECs. **a**) Heatmap showing a selection of differentially expressed genes in CD34^+^CD184^+^CD73^+^DLL4^+^CD43^neg^ (referred to as DLL4^+^, in green) or CD34^+^CD43^neg^CD184^neg^CD73^neg^CD32^+^ (referred to as CD32^+^, in orange) samples isolated at day 8 of WNTd hPSC-derived hematopoietic culture in *n*=3 independent experiments (#1 in light blue, #2 in purple, #3 in yellow) performed in H1 hESCs. Scaled rlog gene expression values are shown in rows. Tiles referring to differentially expressed genes are coloured according to up- (red) or down- (blue) regulation; **b**) Scorecard dot plot showing landmark genes as reported in ^40^. The differential expression was evaluated in *CDH5^+^RUNX1^+^PTPRC^neg^* cells that either express *FCGR2B* (*FCGR2B*>0) or not (*FCGR2B*=0); **c**) UMAP visualization of manually annotated cells and colour-coded by cell type. H1 hESCs, *n*=1. HECs: hemogenic endothelial cells; ECs: endothelial cells; EndoMT: endothelial-to-mesenchyme transition; M phase: mitotic phase; **d**) Feature plot showing *RUNX1* expression across clusters of single-cell data set in Fig. 4a; **e**) Feature plot showing *FCGR2B* expression across clusters 0, 1, 2, 11, 16, 17 that display *RUNX1* as a differential marker; **f**) UMAP visualization of the pseudotime analysis by monocle3 showing the principal graph nodes and trajectory on the clusters with differential expression of RUNX1 as obtained by scRNA sequencing and shown in Fig. 4a, b and Extended Data Fig. 4c. Cells are coloured according to pseudotime; **g**) Pseudotime kinetics of the expression alteration of selected genes (*H19, KCNK17, RUNX1, MYB, SPN*) along the clusters with differential expression of *RUNX1* (clusters 0, 1, 2, 11, 16, 17) as shown in Fig. 4a, Extended Data Fig. 4c, d. Cells are coloured by the cluster identity. Lines denote relative average expression of each gene in pseudotime.

